# Hyperacuity Bayesian methods to enhance temporal resolution of two-photon recording of the complex spikes in the cerebellar Purkinje cells

**DOI:** 10.1101/220350

**Authors:** Huu Hoang, Masa-aki Sato, Mitsuo Kawato, Keisuke Toyama

## Abstract

Two-photon imaging is a major recording technique in neuroscience but its low sampling rate imposes a severe limit of elucidating high temporal profiles of neuronal dynamics. Here we developed two hyperacuity Bayesian algorithms to improve spike detection and spike time precision, minimizing the estimation error supervised by the ground-truth given as the electrical spike signals. The benchmark showed that our algorithms outperformed other unsupervised algorithms maximizing the likelihood of the estimates for both experimental and simulation data. We argue that the supervised algorithms are useful tools to improve spike estimation of two-photon recording in case ground truth signals are available.

## 1 Introduction

Recently, two-photon imaging has been one of major recording means of multi-neuronal activities in neuroscience [Buzsáki, 2004, Denk et al., 1994, Stosiek et al., 2003]. It has high spatial resolution to inform of precise morphology and location of the target neurons. The disadvantages of this technique are low temporal resolution due to serial optical scanning imposing a trade-off between the number of recorded neurons and temporal resolution and the noisiness of optical signals detected by the two photon recording [Hamel et al., 2015, Harris et al., 2016, Katona et al., 2012]. The key techniques for two-photon recording are spike detection and spike time estimation for sparse and noisy spike signals sampled by relatively low frame rate [Grewe et al., 2010, Ji et al., 2016].

A number of studies attempted to resolve the difficulty of spike detection for the noisy and sparse spike signals, including conventional thresholding [Tsutsumi et al., 2015], de-convolution [Yaksi and Friedrich, 2006], template matching [Greenberg et al., 2008], Bayes inference [Deneux et al., 2016, Pnevmatikakis et al., 2016, Vogelstein et al., 2012] and machine learning [Sasaki et al., 2008, Theis et al., 2016]. Rather few studies aimed to improve both spike detection and spike time estimation of two-photon recording. These unsupervised algorithms assumed forward models for spike generation and sampling of spike signals and maximized the likelihood of the estimates of the model parameters [Grewe et al., 2010, Oñativia et al., 2013, Pnevmatikakis et al., 2016].

Here we report two novel algorithms to improve spike estimation of two-photon recording by supervised approach minimizing the estimation error based on the ground-truth given as the electrical spikes. Hyperacuity support vector machine (HSVM) combines Bayesian inference to estimate the most likely spike model with SVM for spike detection using the ground truth signals as teaching signals and for spike time estimation minimizing the sum of squared residuals of spike model predictions for signals. Hyperacuity Bayes (HB) adopts Bayesian hierarchical spike state model and estimate spike states and model parameters by expectation maximization (EM) algorithm with hyperacuity model search. The performance benchmark to compare the foregoing algorithms with ours confirmed superiority of supervised over unsupervised algorithms.

## 2 Methods

### 2.1 Benchmark data

The benchmark data consisted of experimental (EXP) and simulation (SIM) of Ca^2+^ responses. EXP data were simultaneous recordings of electrical (sampling rate, 10 KHz) and two-photon recording (7.8 Hz) of thirty-six C- spikes in five P- cells sampled from five mice injected with Cal-520 dye (9, 6, 5, 5 and 11 C- spikes sampled from #1, 2, 3, 4 and 5 cell, respectively) [Tsutsumi et al., 2015] We conducted a leave-one-out cross-validation for EXP data using the data of four cells for training and those of one cell for test.

We conducted simulation of Ca^2+^ responses for C- spikes with slow [Tsutsumi et al., 2015] and fast dynamics [Blenkinsop and Lang, 2006, Lang, 2002] (Figure 1A). Spike events were generated according to Poisson distribution whose mean firing rate and refractoriness (0.2 Hz, and 1 s, and 1Hz and 0.1s for slow and fast dynamics) were selected so as to minimize KL divergence between SIM and EXP inter-spike interval (ISI) histograms (0.32 and 0.21, Figure 1B). The spike events were convolved with double exponentials (*τ*_1_ = 0.05 s, *τ*_2_ = 0.4 s, see below) whose parameters were estimated from EXP data by Bayesian inference (Figure 1A) and Gaussian noise was added so as to reproduce the SNR (10) of the EXP data. A total of five hundred spikes were generated, 250 each for training and test for all algorithms.

**Figure 1:**
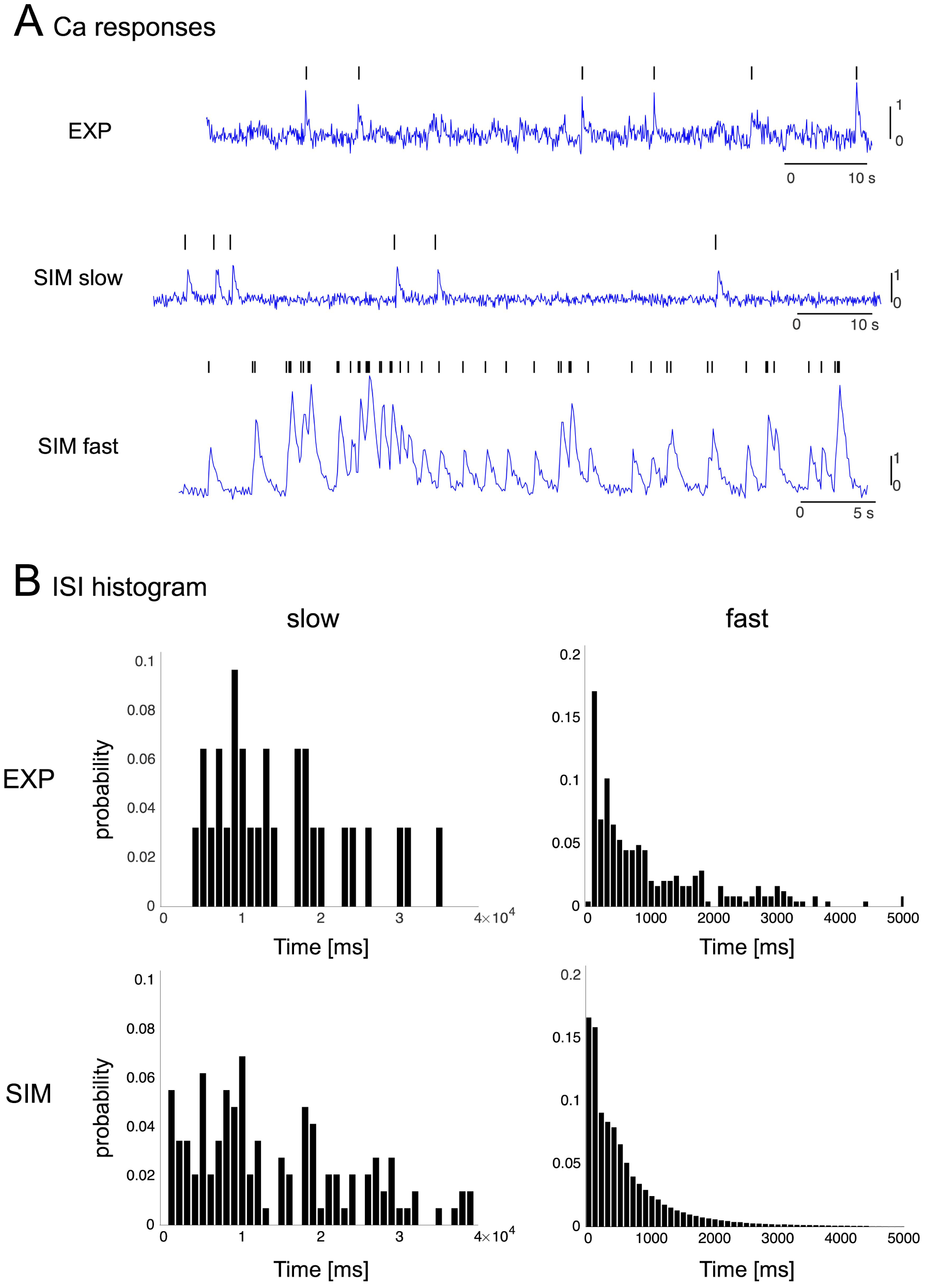
Ca^2+^ responses and ISI histograms of experimental and simulation data. A: Representative Ca^2+^ response of experimental (top, [Tsutsumi et al., 2015]) and simulation data with slow (middle) and fast (bottom dynamics. Dark bars indicate the spike times of simulation data. B: ISI histograms of the experimental (left) and simulated (right) spike data for slow (upper panel) and fast (lower panel) dynamics.

### 2.2 Hyperacuity SVM (HSVM)

#### 2.2.1 Bayesian estimation of the spike model for training set of EXP data

We sampled thirty-six Ca^2+^ spike segments containing the electrical spikes from the training data for EXP data, and estimated the spike model by Bayesian inference assuming that all Ca^2+^ responses originate from the spike model *g*(*t, T, τ*) but are variable due to the noise and sampling jitters [Grewe et al., 2010, Lütcke et al., 2013]:

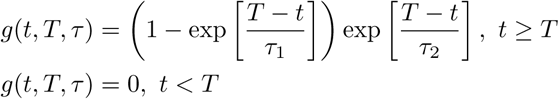
where *t*, *T*, *τ* = (*τ*_1_*, τ*_2_) are the time course, the spike onset, rise and decay time constants, respectively. The time constants were estimated by maximizing the likelihood with a grid-search algorithm:

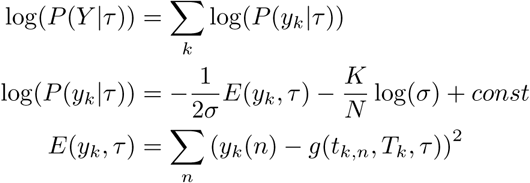
where *Y* = [*y_k_|k* = 1 : *K*] is a set of *K* spike segments (*K* = 36) each of which has *N* (*N* = 16) data points sampled around the electrical spikes, *σ* is the variance of Gaussian noise and *T_k_* is the time of the electrical spike for the *k*-th spike data segment after the onset of the segment.

#### 2.2.2 Supervised learning of SVM with training data

We evaluated the coincidence scores as convolution of the first-derivative of the spike data (*s*) with that of the spike model (*g*) across the entire spike data, *ds/dt ∗ dg/dt*, and sampled the data segment that exceeded the threshold (0.5 SD), including one and seven data points before and after the time for the coincidence score to exceed the threshold (cf. Figure 3). SVM was trained to classify the sampled data segments into spike or non-spike segments using the sampled data and the coincidence scores as the primary and attribute inputs, and the electrical spikes as the teaching (ground truth) signals, respectively.

#### 2.2.3 Spike detection and spike time estimation by HSVM for test data

The trained SVM was used to detect the spike segments sampled from the test data. The pseudo-spike time was estimated as the time when the coincidence score exceeded the threshold. For spike time estimation of detected spikes, we assumed that the spike signals were sampled from the spike model with sampling jitter (difference between pseudo- and true spike time), and estimated the sampling jitter so as to minimize the residuals between the signals and the prediction of spike model by systematically changing the jitter values using ×10 fold finer time bins than the sampling interval (hyperacuity setting). The true spiking time of spike segments was calculated as sum of the pseudo-spike time and the sampling jitter. We subtracted the trace of the preceding spike from the succeeding one for spike time estimation for cases where spikes occurred at short intervals.

### 2.3 Hyperacuity Bayes (HB)

HB consists of the trained and untrained versions. The trained version utilizes ground-truth signals given as the electrical spikes for threshold and spike model optimization, whereas the untrained version does not use such information. We estimated the spike model whose parameters are the spike amplitude (*a*), time course (time constants of double exponentials *τ*), biases (*b*, *b*_0_) and noise (*σ*) using the expectation maximization (EM) algorithm by assuming the constancy of the spike model in the shape and the amplitude. Data segments were sampled by thresholding and the spike states, e.g., the number of spikes contained in the segments and onset times of the individual spikes, were estimated by hyper-acuity Bayesian inference for all data segments so as to maximize the marginal likelihood. Assumed data structures and probabilistic models are described in the first section and detailed procedures of the two versions of HB in the next two sections.

#### 2.3.1 Data structures and probabilistic model

Let us suppose that *K* data segments were sampled from data by thresholding, while leaving the rest data (*y_rest_*).

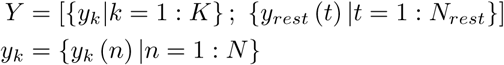
where *y_k_* (*n*) is sampled data at sampling time *t_k,n_* = (*n −* 1) *dt*_0_ + *t_k,_*_1_ of the *k*-th window, *dt*_0_ is sampling step of the observed data with the sampling frequency *f*_0_ = 1*/dt*_0_.

Probabilistic model for spike states is given by

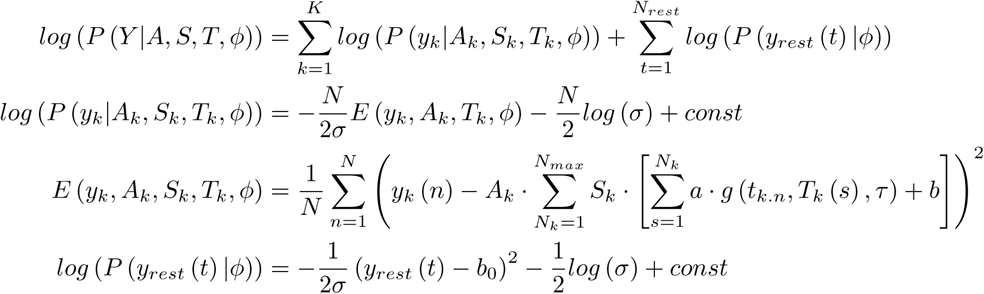
where *A_k_* is a spike indicator variable, which represents presence or absence of spikes in *k*-th window, and takes binary value (0 or 1). *S_k_* represents a spike state in *k*-th window and takes binary vector value (Potts spin variable)

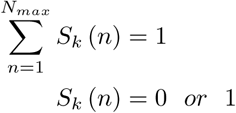

*S_k_* (*N_k_*) = 1 means that there are *N_k_* spikes in k-th window. Maximum number of spikes in a window is assumed as *N_max_*. *T_k_* is a set of spike onset times in *k*-th window. *T_k_* (*s*) is *s*-th set of spike onset time configuration and was estimated with shorter time step *dt* (hyper-acuity setting: *dt < dt*_0_). *a* represents the amplitude of spike response function. *b* and *b*_0_ represent bias in spike and no-spike regions, respectively. *σ* is the variance of Gaussian noise. A set of global parameters are denoted as *ϕ* = (*a, τ, b, b*_0_*, σ*) and is assumed to be common for all spikes and the rest data.

We assume hierarchical non-informative prior

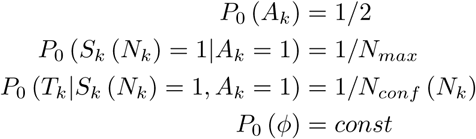

where, *N*_*conf*_ (*N*_*k*_) represents number of configurations of *T*_*k*_ for *N*_*k*_ spike case.

#### 2.3.2 Spike estimation with training data for the trained version

- Sampling of data segments We first extracted continuous regions in which the signals exceed the threshold. Next, each region is segmented into fixed-length data segments of 8 points. We divided two overlapped data segments, whose onset intervals were less than the length of data segment (8 points), into three divided non-overlap segments. For examples, two segments of points #1-10 and #5-14 were divided into three non-overlap segments of points #1-4, #5-10 and #11-14, respectively.
- Threshold optimization Optimization of threshold was conducted so as to maximize true positive cases and minimize the false positive cases referring to the ground-truth given by the electrical spikes.
- Estimation of spike model parameters for training data The spike model parameters including spike amplitude (*a*), biases (*b*, *b*_0_) and noise variance *σ* were estimated by the EM algorithm (see Appendix for EM algorithm and posterior probability), while time constants of spike response (*τ*) was estimated by iterative alternate coordinate 1D grid search, because the log-likelihood surface w.r.t. *τ* is highly non-linear and its curvature is positive indefinite.
- Maximum number of spikes per data segments The maximum number of spikes contained in single data segments was estimated for the 95-percentile value of the spike number histogram of the training data segments.
- Estimation of posterior probability for training data Posterior probability of the spike states were estimated for each data segment assuming that they are independent from each other. For overlapped segments, posterior probabilities were integrated among the overlapped ones by Bayes inference.
- Training of multi-nominal classifier with training data We used a multi-nominal classifier to predict the number of spikes from posterior probability *P* (*A_k_* = 1*, S_k_* (*N_k_*) = 1*|y_k_, ϕ*) *, N_k_* = 1 : *N_max_*.
- Estimation of spike number and spike onset time for test data The data segments were sampled for the test data by thresholding whose threshold was optimized for the training data, and posterior probability of spike state for each segment is calculated in the same way for the training data. The number of spikes was estimated from posterior probability for number of spikes, *P* (*A_k_* = 1*, S_k_* (*N_k_*) = 1*|y_k_, ϕ*) *, N_k_* = 1 : *N_max_*, using the multi-nominal classifier trained for the training data, and spike onset times were finally estimated with hyper-acuity time step by maximising log-likelihood for the estimated number of spikes. The contributions of preceding spikes were subtracted from the data signal as was done for HSVM.

#### 2.3.3 Spike estimation by untrained version of HB

The untrained version of HB was different from the trained version in two ways. First, to obtain initial estimate of spike states, we assumed each data segment contains only one spike and the peak of data segment corresponds to the peak of spike response curve. We varied threshold value and find the optimal one that gives maximum log-likelihood. This initialization process was done for each *τ* value in the 1D grid search. Starting from the initial estimate, iterative alternate coordinate 1D grid search with EM algorithm was conducted. Second, the number of spikes was determined for each data segment which gives the maximum posterior probability for given number of spikes 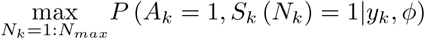.

### 2.4 Comparison of performance with the three benchmark algorithms

We evaluated the performance for the SIM and EXP data for the three foregoing algorithms, including the peeling algorithm [Grewe et al., 2010], the finite-rate innovation method (FRI) [Oñativia et al., 2013] and the Monte-Carlo Markov Chain method (MCMC) [Pnevmatikakis et al., 2016] in the untrained and trained conditions.

### 2.5 Performance analysis

Correct hit, false alarm and false negative cases were defined as the cases where the time intervals of the reconstructed and true spikes were ≤ 0.15s, those of the reconstructed spikes from the true spikes and those of the true spikes from the reconstructed spikes were > 0.15 s, respectively. ROC analysis was conducted as:

- Sensitivity = Hit / (Hit + False negative)
- Precision = Hit / (Hit + False alarm)
- F1-score = 2 × (Sensitivity × Precision) / (Sensitivity + Precision)

For evaluation of spike time accuracy, spike time errors were estimated as the absolute time difference between true and reconstructed spikes for correct hit cases.

## 3 Results

### 3.1 HSVM

#### 3.1.1 Bayes estimation of spike model

We estimated the spike model by Bayesian inference for the thirty-six data segments (thin dark traces in Figure 2) that included both electrical and two photon recording of Ca^2+^ responses, assuming that the Ca^2+^ responses were sampled from the spike model ((1−exp(−*t*/*τ*_1_) exp(−*t*/*τ*_2_)) with sampling jitters, superimposed of Gaussian noise. In agreement of this view, the spike model (*τ*_1_ = 0.05, *τ*_2_ = 0.4s, red trace) was slightly larger in amplitude and faster in rise and decay time than the electrical spike-triggered average of the Ca^2+^ response (blue trace) probably due to temporal dispersion of the sampling.

**Figure 2:**
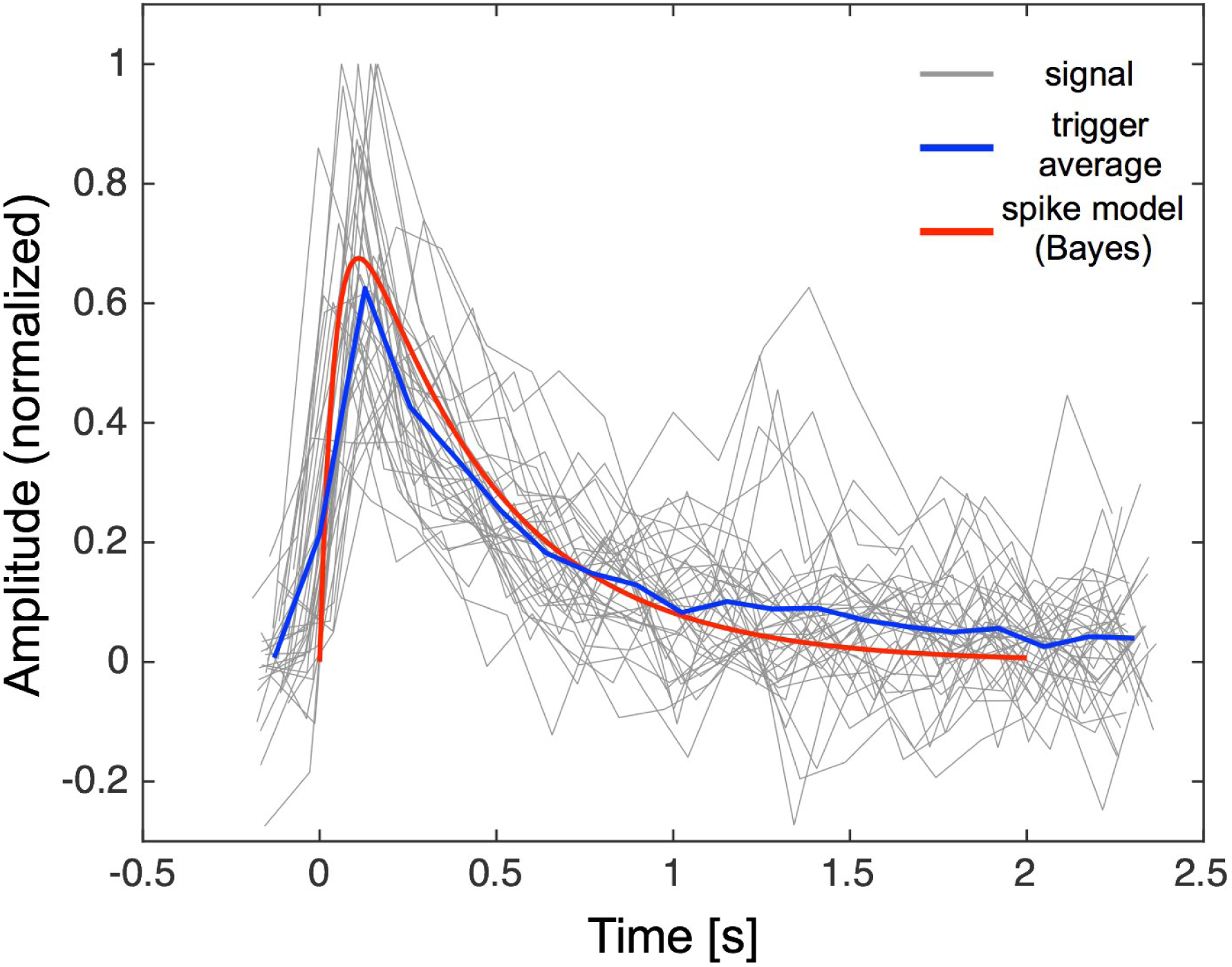
Bayesian estimation of spike model (red trace). Grey traces represent individual Ca^2+^ responses of five P- cells whose onsets are synchronized with those of electrical spikes. Blue and red traces represent spike-triggered average for the electrical spikes and Bayesian estimation of Ca^2+^ responses, respectively.

#### 3.1.2 Spike detection

We estimated the coincidence score between the first order derivative of the Ca^2+^ signals and the spike model (*ds/dt ∗ dg/dt*) and sampled the data segments as the candidates for the spikes at the point when the coincidence score exceeded the threshold (0.5 SD, dashed dark line in Figure 3A-C). Coincidence thresholding (open dark circles in Figure 3A-C) detected true (short grey bars) as well as many false alarms spikes. By contrast, SVM selected true spikes (filled dark circles) rejecting false alarm spikes from the spike candidates that contained many false alarm spikes as well as true ones.

**Figure 3:**
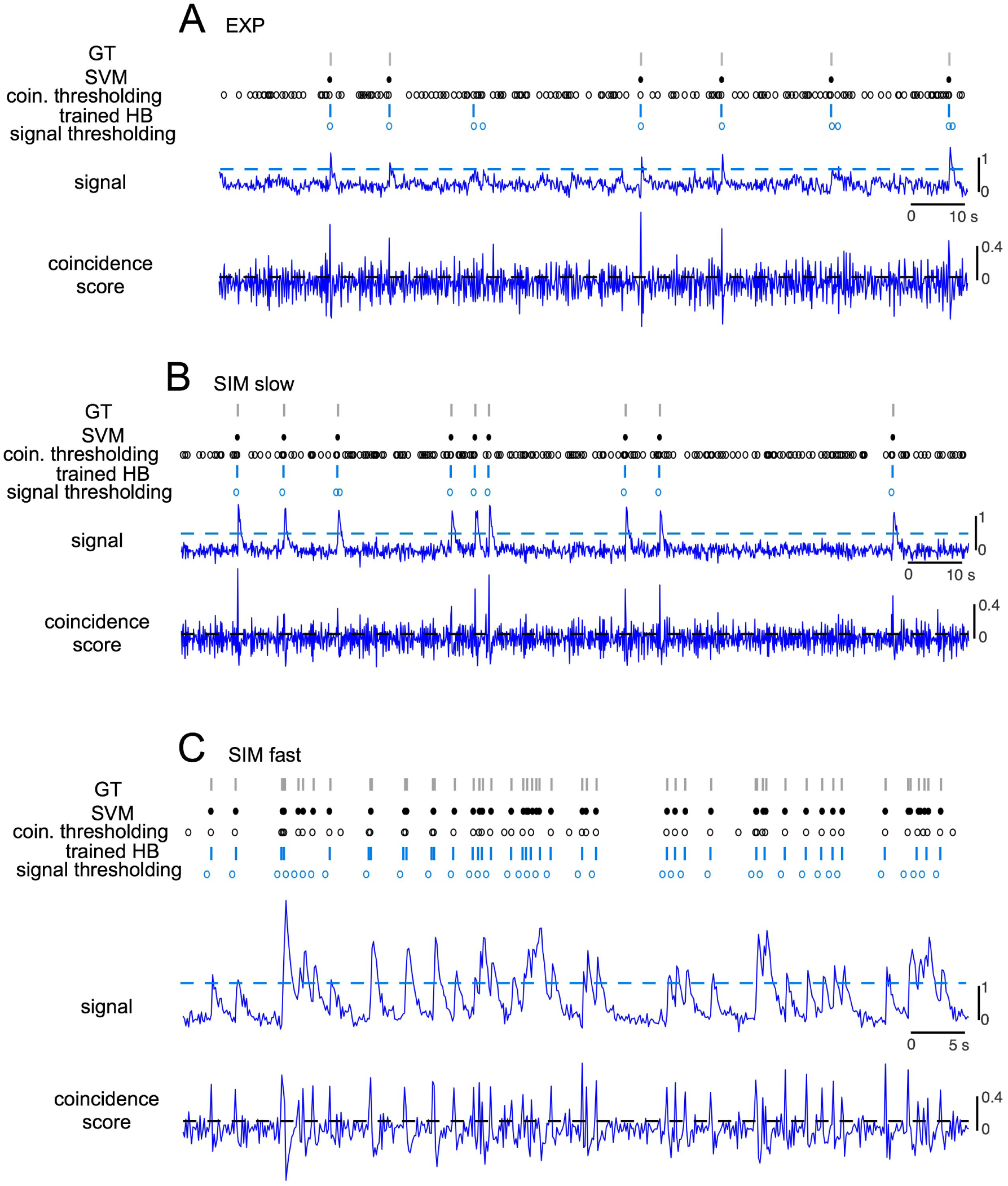
Spike detection by HSVM and HB. A: Ca^2+^ responses, coincidence scores and spike detection by thresholding of coincidence scores (open dark circles), spike detection by SVM (filled dark circles), thresholding of raw signal by HB (open blue circles), spike estimation by HB (blue bars) and the ground truth (grey bars) for experimental data. The dashed dark and dashed blue lines indicate 0.5SD of coincidence scores and the optimized threshold by HB, respectively. B and C: those for SIM data with slow and fast dynamics.

We tentatively assumed that the spike time of the detected spikes as the time when the coincidence exceeded the threshold (pseudo-spike time) and estimated sampling jitters of the pseudo spike time to the true spike onset by Bayesian inference using hyperacuity vernier (10-fold finer time bin than the sampling interval) and the true spike times were calculated as sum of pseudo-spike time and sampling jitter (dark bars in Figure 4A-C). Figure 4 exemplifies the performance of HSVM for EXP data and SIM data with slow and fast spike dynamics. Comparison of spikes detected by HSVM (dark bars) with the ground-truth (electrical spikes, grey bars) indicates that HSVM almost perfectly detected spikes and correctly estimated the spike time across the three cases.

**Figure 4:**
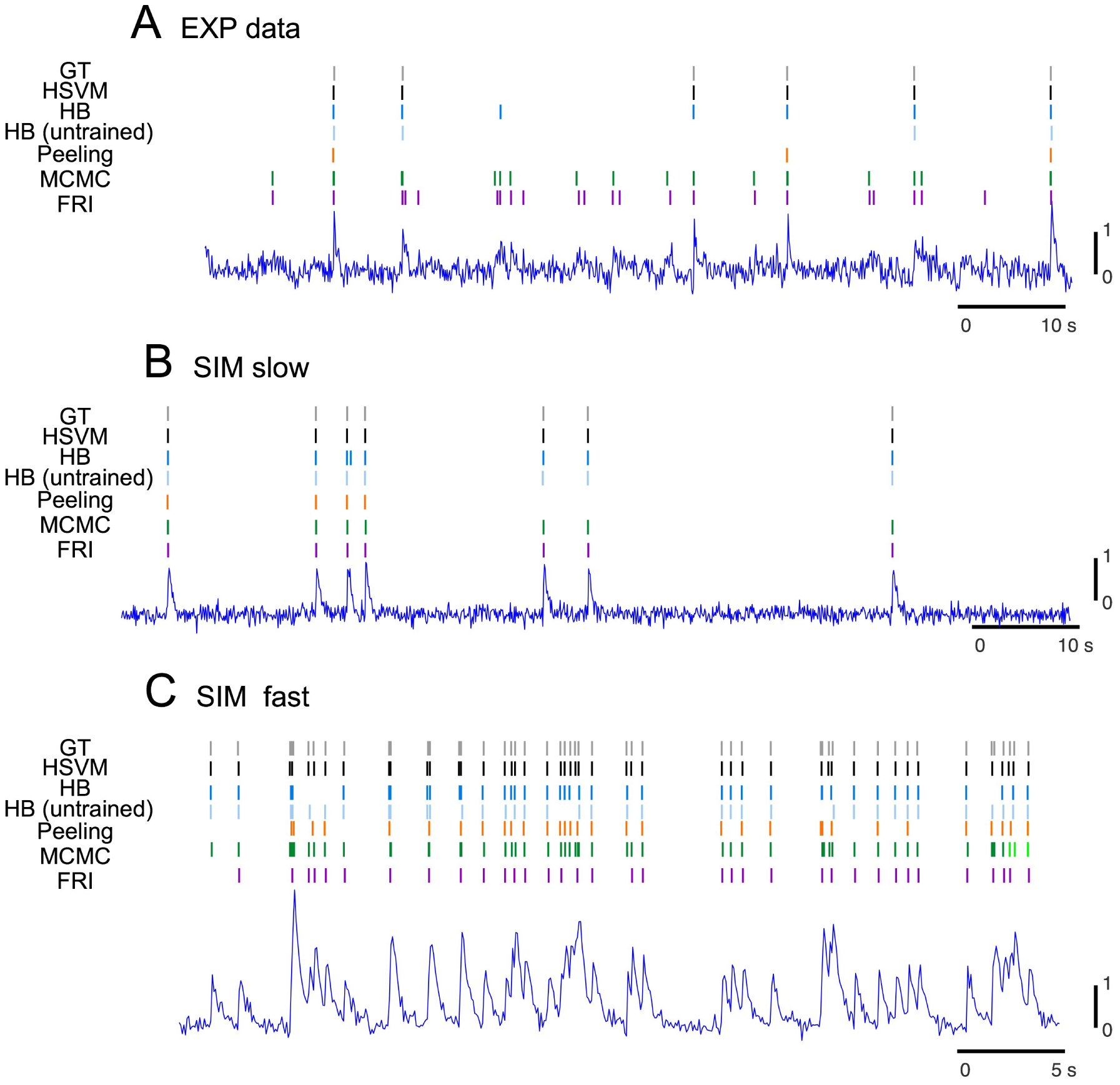
Comparison of spike detection by HSVM and HB with those for the three other algorithms. A, B and C: spike detection is shown for experimental and simulation data with slow and fast dynamics, respectively, as short bars of different colors for Ca^2+^ responses (bottom traces).

Next, we conducted ROC analysis to evaluate performance of spike detection of the HSVM for training and test SIM data, each including 250 spikes as well as training and test EXP data including 144 and 36 spikes shuffled by 5-fold-cross-validation. The performance of HSVM evaluated as sensitivity, precision and F1 score was perfect for slow SIM data (100, 100, and 100%, dark columns in Figure 5A-C) but slightly reduced for fast SIM data (97, 91, and 94%) as well as EXP data (80, 97, and 87%).

**Figure 5:**
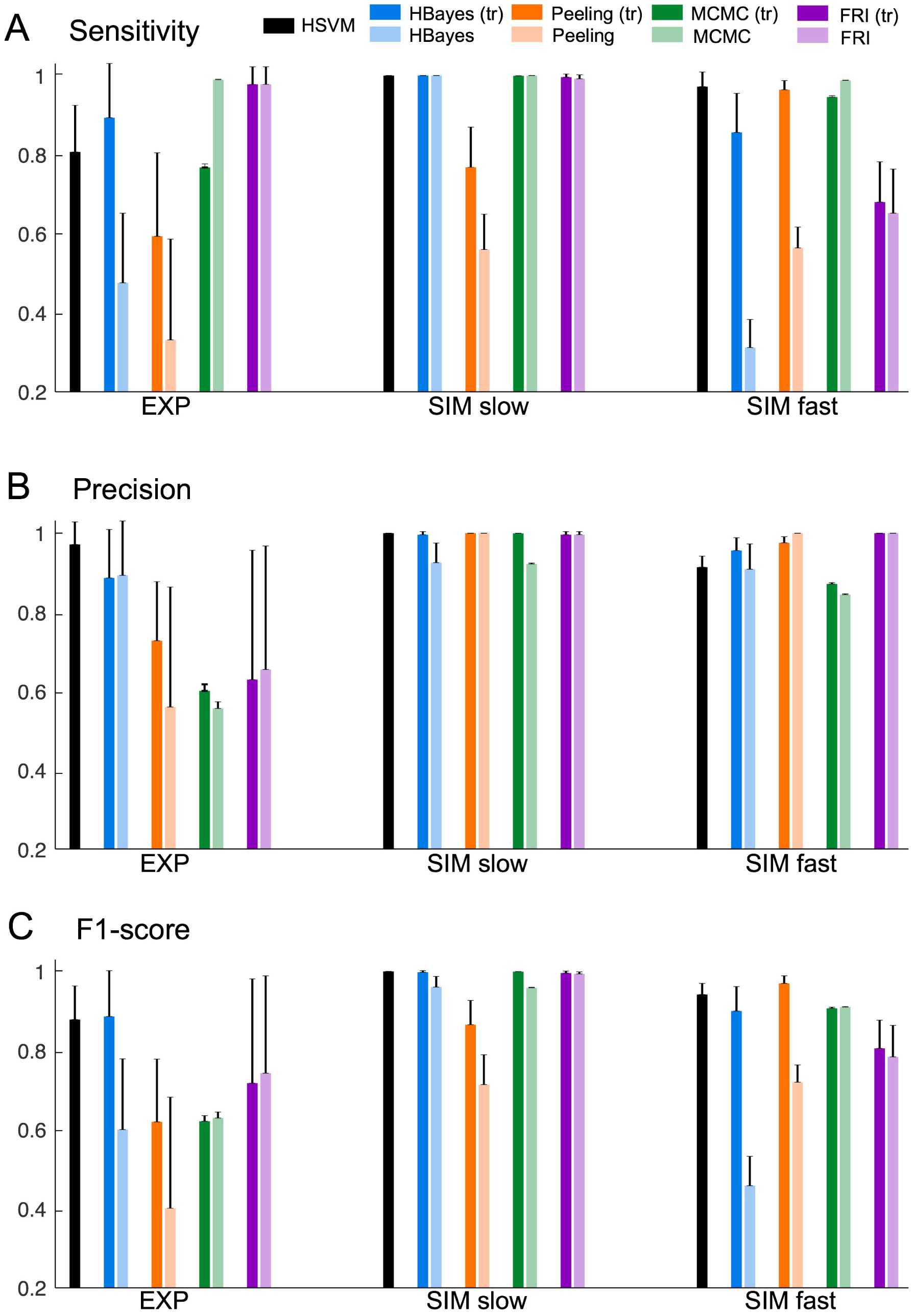
ROC analysis of spike detection for five algorithms. A, B and C, sensitivity, precision and F1 score for HSVM (dark columns), HB (blue columns), Peeling algorithm (orange columns), MCMC (green columns) and FRI (purple columns). The dark and light conventions represent the scores for trained and untrained versions for HB and those (Peeling, MCMC, and FRI) for the spike model parameters estimated by our Bayesian inference and those listed in the papers.

#### 3.1.3 Spike time estimation

Performance of HSVM to estimate spike time was evaluated as the spike time errors (the time difference between true and reconstructed spikes) for correct hit cases. HSVM detected 100, 97 and 80% of the spike with spike time errors of 12 ± 13 ms, 20 ± 19 ms and 40 ± 35 ms for the SIM slow and fast dynamics and EXP data, respectively (Figure 6, dark columns). The hyperacuity performance of HSVM, evaluated as the ratio of the sampling interval to the mean of spike time errors, was 8, 5 and 3.2 folds for the SIM slow and fast dynamics (sampling interval, 100 ms) and EXP data (128 ms), respectively.

**Figure 6:**
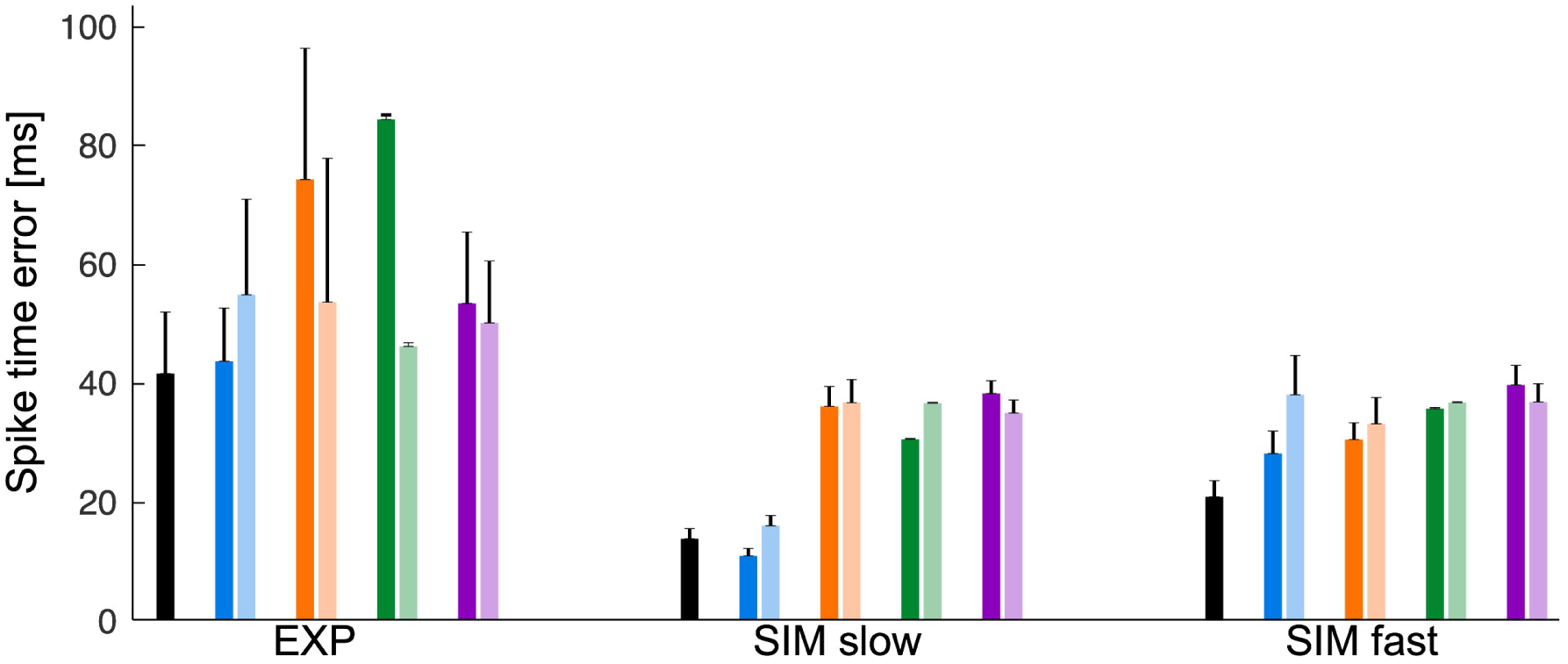
Spike time errors for EXP and slow and fast SIM data. Figure conventions are same as those in Figure 5.

### 3.2 Hyperacuity Bayes (HB)

The trained version of HB was conducted in two steps. First, spike data segments were sampled by thresholding (Figure 3A-C, dashed blue lines) whose threshold was optimized so as to maximize the correct hit and minimizing the false alarm, and the spike model was estimated for the sampled data segments by EM algorithm (blue open circles). Second, the number of spikes and the spike times were estimated by the multi-nominal classifier and Bayesian inference, respectively (blue bars).

The trained version of HB detected all spikes for EXP data (Figure 4A, dark blue bars) and slow SIM data (Figure 4B) but missed a few spikes for fast SIM data (Figure 4C), while untrained version missed significant number of spikes in fast SIM data and EXP data (Figure 4A-C, light blue bars). ROC analysis confirmed that trained version performed perfectly for slow SIM data (sensitivity, precision, and F1 score, 100, 100, 100%, Figure 5A-C, dark blue columns) and with a good accuracy for EXP data (89, 88, 88%) as well as fast SIM data (85, 95, 90%). The untrained version performed rather poorly for all of EXP, slow and fast SIM data (sensitivity: 47, 100, 31%, precision: 89, 92, 90%, F1-score: 60, 96, 45%, Figure 5A-C, light blue columns). Altogether, performance of the trained version of HB was as good as that of HSVM for all types of data. Correspondingly, the spike time errors of the trained version of HB were slightly smaller than those of the HSVM for slow SIM data but larger for fast SIM data and EXP data, with hyperacuity performance of 10, 3.5, 3 folds (Figure 6, dark and light blue columns).

### 3.3 ROC and spike time error analysis for the three other algorithms

We conducted ROC and spike time analysis of the three other algorithms including Peeling, MCMC and FRI algorithms as benchmarks for performance of our algorithms. Sensitivity for MCMC was as high as for HSVM in all of the three data (cf. Figure 5, light green columns) for SIM slow (100%) and fast dynamics (98%) and EXP data (98%), and that for FRI was also high for slow SIM data (99%) and EXP data (97%) (light purple columns), but relatively low (65%) for fast SIM data, and that for Peeling was rather poor across all of three data (light orange bars, 33, 56 and 56% for EXP and SIM data with slow and fast dynamics, respectively). Precision of the three algorithms was high for slow SIM data, and that of Peeling and FRI remained also high for fast SIM data (Figure 5B, light orange and purple columns, 100, 100%) but that of MCMC became rather low for fast SIM data as well as for EXP data (light green columns, 84 and 55%). Accordingly, F1 scores for MCMC and FRI were as high as that for HSVM for slow SIM data (Figure 5C, light green and purple columns, 96 and 99%), but that for Peeling was significantly worse than HSVM (71%). Those of all three algorithms for fast SIM data and EXP data were considerably lower (71, 91, 78% and 40, 62, 74% for Peeling, MCMC and FRI, respectively) than those for HSVM. Spike time errors of the three algorithms were all significantly larger (roughly 3 and 2.5 folds for SIM and EXP data, respectively, cf. Figure 6, light orange, green and purple columns) than those of HSVM.

Altogether, HSVM either outperformed or almost equally performed in sensitivity, precision and F1-scores compared with the other three algorithms. Most importantly, HSVM retained significantly higher precision than the other algorithms across all types of spike data, and accordingly showed the highest F1-scores. Performance of spike detection for trained HB was also comparable with that for HSVM except for relatively low sensitivity for fast SIM data. Precision of spike time estimation, evaluated as the spike time errors, was also highest for HSVM and next for trained HB of all algorithms in all of the three types of data.

We finally tested whether the accurate information of spike model parameters such as the amplitude and the time constants of the rise and decay exponentials may improve the performance of the three benchmark algorithms by comparing the performance of those algorithms between the initial values in the papers and those estimated by our Bayesian estimation (cf. light and dark bars in Figures 5 and 6). It was found that the accurate information did not significantly improve the performance of spike detection or spike time estimation of the three benchmark algorithms except for Peeling algorithm for which there was rather significant improvement in F1 score (60, 87 and 92% for EXP and slow and fast SIM data, respectively).

## 4 Discussions

We found that our two algorithms using the supervised approach outperformed the three benchmark algorithms using the unsupervised approach in the spike detection as well as spike time estimation for most of EXP and slow and fast SIM data. Our algorithms were superior to the unsupervised algorithms in F1 score that is the most important measure of spike detection performance in all of EXP, slow and fast SIM data. The superiority of supervised algorithms was also evident in the performance of spike time estimation. The spike time error was minimal for HSVM and next minimal for HB for all of the three types of data.

It is noteworthy that the superiority of the supervised algorithms over the unsupervised algorithms was most pronounced for EXP data in both spike detection and spike time estimation. This finding is in agreement with the recent study [Theis et al., 2016] reporting that the unsupervised algorithms for spike detection perform significantly worse for EXP data than SIM data. The reduced performance of the unsupervised algorithms for EXP data might be due to the factors such as non-Gaussian noise and variations of the spikes in EXP data that are unforeseen by the forward models for the likelihood maximization. Conversely, those effects may be compensated by error feedback in the supervised algorithms. In consistency with this view, the trained version of HB significantly outperformed the untrained version.

In sum the supervised algorithms detected spikes with high F1 score (90%) and estimated spikes with more than 3-folds temporal precision of the frame rate and indicate that they are useful tools to improve spike estimation of two-photon recording in case ground truth signals are available.

## Acknowledgements

We thank Dr. Kazuo Kitamura and Dr. Shinichiro Tsutsumi for their assistance in collection of the simultaneous recordings of Purkinje cells.

## Appendix

### Expectation maximization (EM) algorithm

In this appendix, we described the expectation maximization (EM) algorithm to estimate model parameters as well as computation of the posterior probabilities of the probabislitic model described in the main text.

### Estimation step (E-step)

In the E-step, posterior probability of hidden states are estimated using current estimate of model parameters *ϕ*. To get posterior probability, joint probability is first calculated by combining loglikelihood and prior. Joint probability for a spike state *{A_k_, S_k_, T_k_}* in *k*-th window is given by:

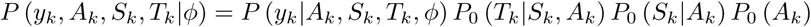

and marginal probability is given by

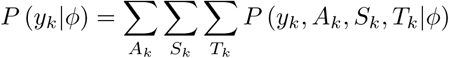

Then posterior probability for a spike state *{A_k_, S_k_, T_k_}* is calcuated as

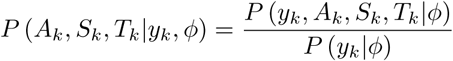

In particular, posterior probability for a given number of spikes *N_k_*, *P* (*A_k_* = 1, *S_k_* (*N_k_*) = 1*|y_k_*, *ϕ*) = ∑*_T_k__ P* (*A_k_* = 1, *S_k_*, (*N_k_*) = 1, *T_k_|y_k_*, *ϕ*), determine the number of spikes in the k-th window. Also, *P* (*A_k_* = 1, *S_k_* (*N_k_*) = 1, *T_k_* (*s*) *|y_k_*, *ϕ*) gives posterior probability for the s-th spike onset time configulation with *N_k_* spikes.

### Maximization step (M-step)

In the M-step, model parameters are updated to new value *ϕ_new_* by maximizing the Q-function defined as

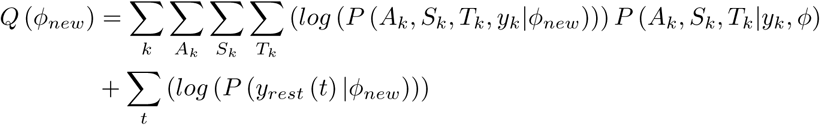

by solving maximum condition

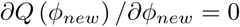

